# BEST: a web application for comprehensive biomarker exploration on large-scale data in solid tumors

**DOI:** 10.1101/2022.10.21.513300

**Authors:** Zaoqu Liu, Long Liu, Siyuan Weng, Hui Xu, Zhe Xing, Yuqing Ren, Xiaoyong Ge, Libo Wang, Chunguang Guo, Lifeng Li, Quan Cheng, Peng Luo, Jian Zhang, Xinwei Han

## Abstract

**Summary:** Data mining from RNA-seq or microarray data has become an essential part of cancer biomarker exploration. Certain existing web servers are valuable and broadly utilized, but the meta-analysis of multiple datasets is absent. Most web servers only contain tumor samples from the TCGA database with only one cohort for each cancer type, which also means that the analysis results mainly derived from a single cohort are thin and unstable. Indeed, consistent performance across multiple independent cohorts is the foundation for an excellent biomarker. Moreover, many analytical functions researchers require remain adequately unmet by these tools. Thus, we introduce BEST (Biomarker Exploration for Solid Tumors), a web application for comprehensive biomarker exploration on large-scale data in solid tumors. BEST includes more than 50,000 samples of 27 cancer types. To ensure the comparability of genes between different sequencing technologies and the legibility of clinical traits, we re-annotated transcriptome data based on the GRCh38 patch 13 sequences and unified the nomenclature of clinical traits. BEST delivers fast and customizable functions, including clinical association, survival analysis, enrichment analysis, cell infiltration, immunomodulator, immunotherapy, candidate agents, and genomic alteration. Together, our web server provides multiple cleaned-up independent datasets and diverse analysis functionalities, helping unleash the value of current data resources. It is freely available at https://rookieutopia.com/.

**The bigger picture:** Bioinformatics web servers enable researchers without computational programming skills to conduct various bioinformatics analyses. However, most web servers only contain tumor samples from the TCGA database with only one cohort for each cancer type, which also means that the analysis results mainly derived from a single cohort are thin and unstable. Thus, we introduce BEST (Biomarker Exploration for Solid Tumors), a web application for comprehensive biomarker exploration on large-scale data in solid tumors. BEST includes more than 50,000 samples of 27 cancer types that have been uniformly re-annotated based on the GRCh38 patch 13 sequences, which ensures the comparability of genes between different sequencing technologies. BEST also offers prevalent functions including clinical association, survival analysis, enrichment analysis, cell infiltration, immunomodulator, immunotherapy, candidate agents, and genomic alteration. Together, BEST provides a curated database and innovative analytical pipelines to explore cancer biomarkers at high resolution.

## Introduction

Biomarker identification is an important goal of cancer research for clinicians and biologists. How to explore specific biomarkers that can distinguish tumoral from normal tissues, identify treatment-resistant patients, predict patient prognosis and recurrence, etc., are routine research tasks. Recently, immunotherapies represented by immune checkpoint inhibitors have opened a new era in cancer treatment, significantly improving the clinical outcomes of cancer patients^1^. However, only a small fraction of patients can generate considerable benefits from immunotherapies^2^. Exploring specific biomarkers that can effectively predict immunotherapeutic efficacy is crucial for preventing under-or over-treatment.

With the advancement of bioinformatics techniques, researchers are inclined to explore cancer biomarkers using RNA-seq or microarray data^3, 4^, and data mining has become an essential part of cancer research. However, these works may be difficult and inconvenient for clinicians and biologists without computational programming skills. Currently, several open-access web servers that allow users to analyze and visualize gene expression online directly are emerging, such as GEPIA^5^, Xena^6^, ExpressionAtlas^7^, and HPA^8^. Although these web applications are valuable and broadly utilized, obtaining high confidence results in a specific tumor is difficult because their data sources are mainly derived from the TCGA database. Consistent performance across multiple independent datasets is the foundation for an excellent biomarker. In addition, the deeper exploration of specific biomarkers on underlying mechanisms, tumor microenvironment, and drug indications are missing in these tools.

To address these unmet needs, we have developed Biomarker Exploration for Solid Tumors (BEST), a web-based application for comprehensive biomarker exploration on large-scale data in solid tumors and delivering fast and customizable functionalities to complement existing tools.

## Results

### Quick start

BEST offers a simple interactive interface. Users first select one cancer type and then determine the input category—single gene or gene list (Figure 1). For the single-gene module, users can enter a gene symbol or an Ensembl ID in the ‘Enter gene name’ field to explore a gene of interest. The gene list module needs users to input a list of genes and pick a method to calculate the gene set score for each sample. The embedded methods include gene set variation analysis (GSVA)^9^, single sample gene set enrichment analysis (ssGSEA)^10^, z-score^11^, pathway-level analysis of gene expression (PLAGE)^12^, and the mean value. Users can customize the name of the gene set score.

**Figure 1.**
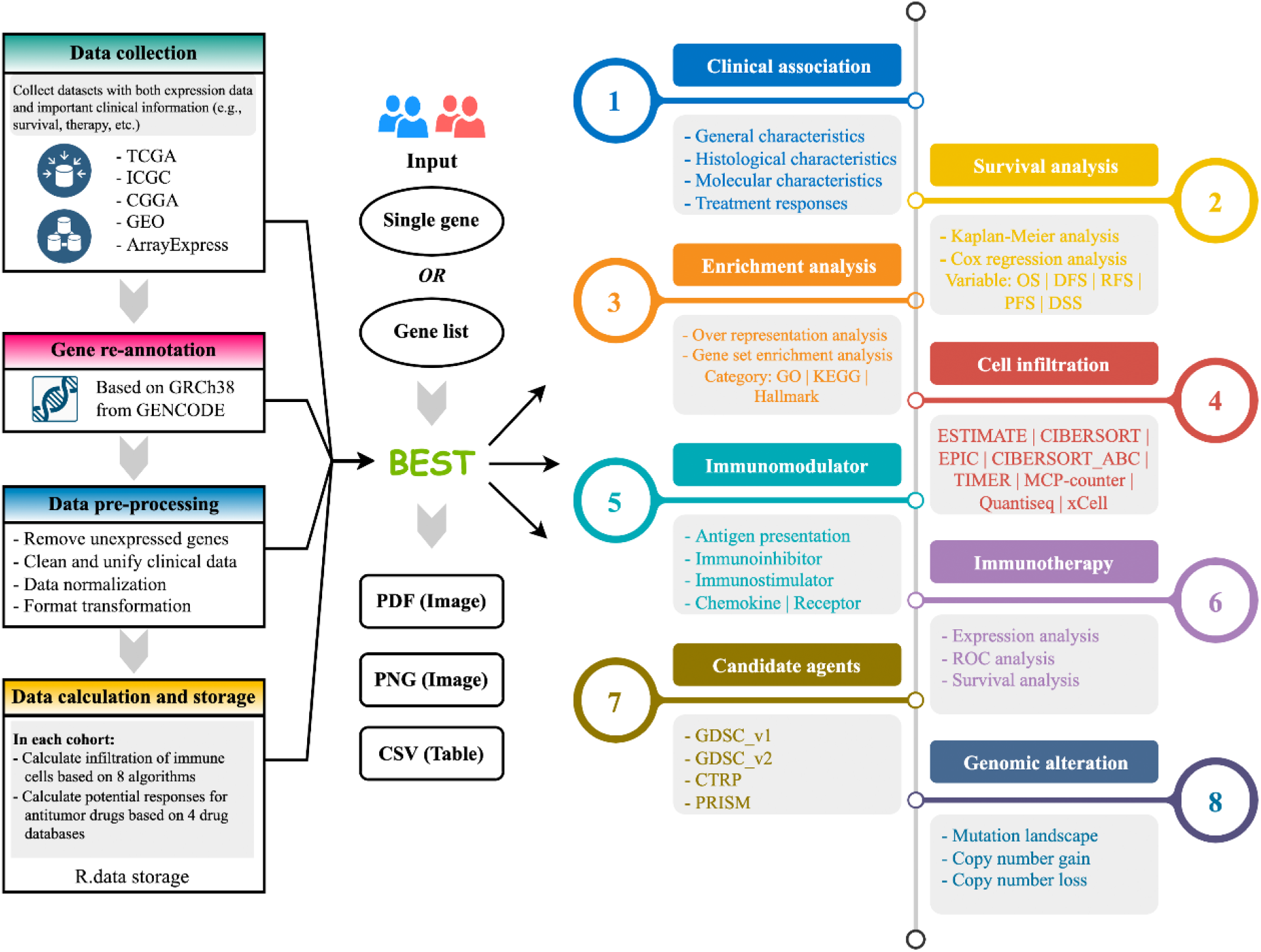
Overview of the BEST analytical framework.

### Clinical association

In this module, users can explore the associations between the expression or score of the input variable and general characteristics (e.g., age, gender, alcohol, smoke, etc.), histological characteristics (e.g., tissue type, tumor site, stage, etc.), molecular characteristics (e.g., TP53 mutation, microsatellite instability, etc.) and treatment responses (e.g., chemotherapy and bevacizumab responses, etc.) (Figure 2A). Whether to use parametric or nonparametric statistical tests for group comparisons based on the distribution of input variable. For example, users can easily explore the differential expression of the input variable between tumor and normal tissues or find the associations between the input variable with smoke and alcohol. Our datasets also include abundant treatment responses, which might contribute to developing promising biomarkers in clinical settings. Importantly, analysis results tend to be displayed in multiple independent cohorts, which provides a reference for the stability power of a variable of interest. For instance, Figure 2B illustrates that CRC tumors process a significantly higher expression of *COL1A2* than normal tissues in most CRC datasets with tissue type information.

**Figure 2.**
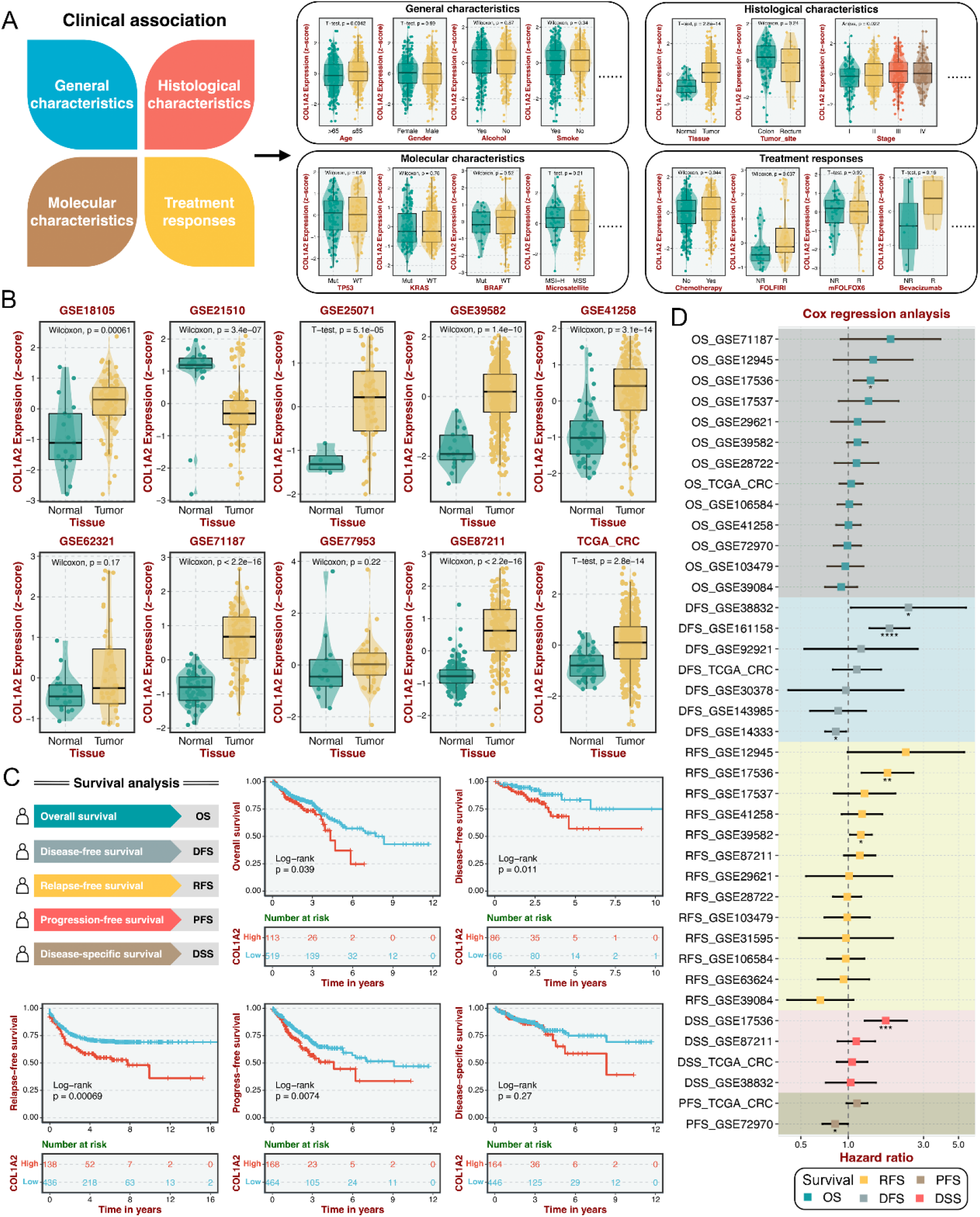
Modules for clinical association and survival analysis. **A**. Four categories of clinicopathologic information are mainly included in the clinical association module. **B**. An example illustrates the differential expression of *COL1A2* in multiple CRC datasets between the tumor and normal groups. **C**. Five categories of survival variables are utilized in the survival analysis module, and examples of five survival variables for Kaplan-Meier analysis. **D**. An example displays the cox regression analysis of five survival variables in multiple CRC datasets.

### Survival analysis

BEST performs survival analysis based on gene expression or gene set score. This module allows users to explore the prognostic significance for overall survival (OS), disease-free survival (DFS), relapse-free survival (RFS), progression-free survival (PFS), and disease-specific survival (DSS) (Figure 2C). BEST generates Kaplan-Meier curves with log-rank test and forest plot with cox proportional hazard ratio and the 95% confidence interval information for various survival outcomes in multiple independent datasets (Figure 2C-D). Kaplan-Meier analysis requires categorical variables, we thus provide five cutoff options for users to choose from, including ‘median’, ‘mean’, ‘quantile’, ‘optimal’, and ‘custom’. For example, when investigating gene *COL1A2* in survival analysis of CRC, users can obtain Kaplan-Meier curves with a specific cutoff approach and a Cox forest plot for five survival outcomes across all CRC datasets with survival information.

### Enrichment analysis

BEST provides two enrichment frameworks: over-representation analysis (ORA) and gene set enrichment analysis (GSEA). Users can select the top gene (self-defined number) most associated with the input variable to perform ORA and apply a ranked gene list based on the correlation between all genes and the input variable to carry out GSEA (Figure 3A). Of note, the final correlation coefficient between the input variable and each gene is the average correlation of all datasets in specific cancer. The output forms of ORA are GO and KEGG bar charts (Figure 3B-C). Also, GSEA results are exhibited using GSEA-Plot (Figure 3D) and Ridge-Plot (Figure 3E) images.

**Figure 3.**
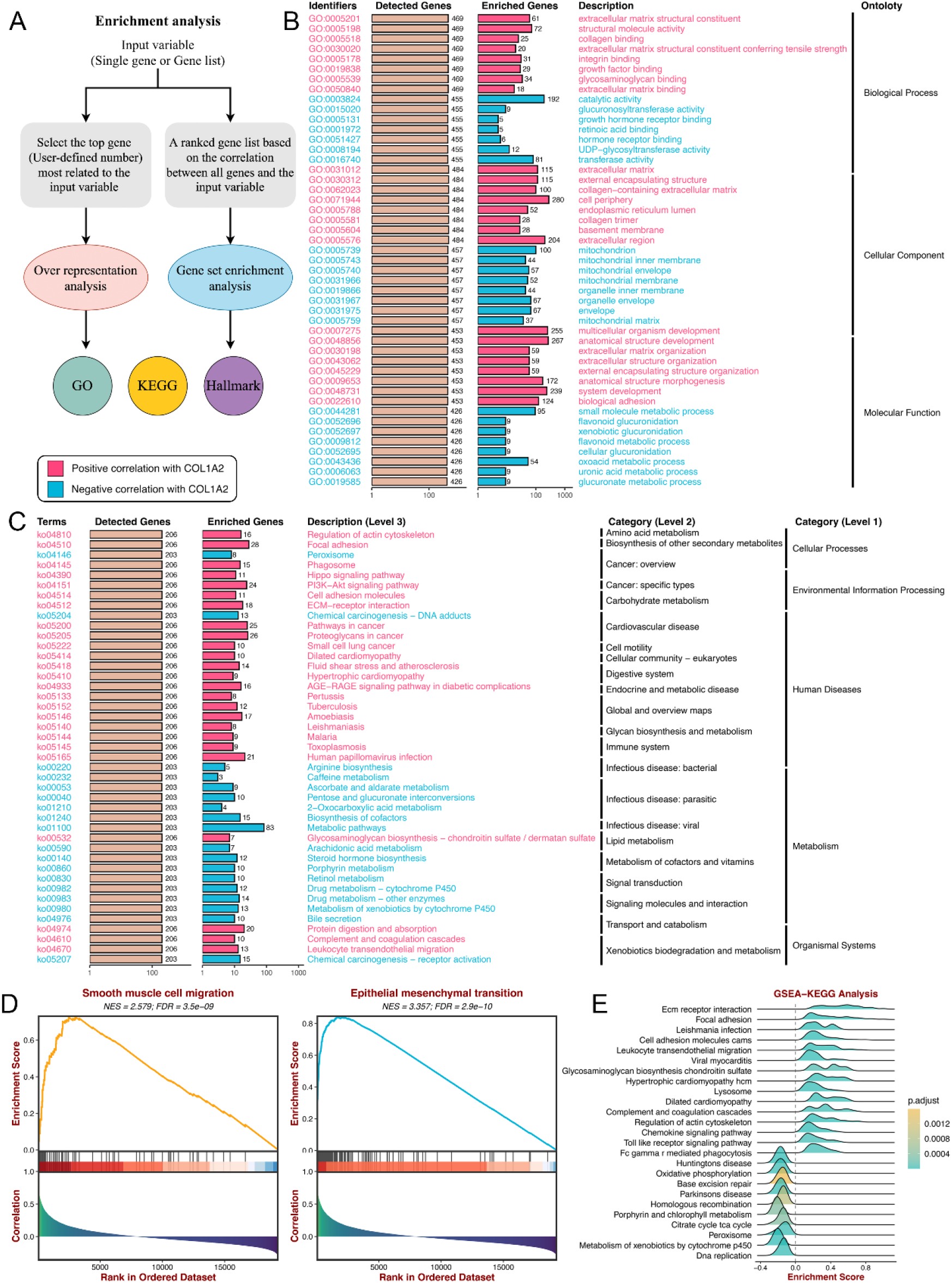
Enrichment analysis module. **A**. Enrichment analysis module includes two enrichment frameworks: over-representation analysis (ORA) and gene set enrichment analysis (GSEA). **B-C**. GO (**B**) and KEGG (**C**) bar charts for the ORA framework. **D-E**. GSEA-Plot (**D**) and Ridge-Plot (**E**) examples for the GSEA framework.

### Cell infiltration and immunomodulator

BEST offers eight prevalent algorithms to estimate immune cell infiltration in the tumor microenvironment (TME) (Figure 4A), including CIBERSORT^13^, CIBERSORT ABS^13^, EPIC^14^, ESTIMATE^15^, MCP-counter^16^, Quantiseq^17^, TIMER^18^, and xCell^19^. To avoid time-consuming calculations for users and save computing resources, these eight algorithms have been executed in advance across all datasets, and the resulting data have been stored in the website background. Additionally, BEST provides five immunomodulator categories: antigen presentation, immunoinhibitors, immunostimulators, chemokines, and receptors (Figure 4A). Users can generate heatmap and correlation scatter plots from these two analysis modules. The heatmaps illustrate the correlations of the input variable with each immune cell/immunomodulator across all cohorts (Figure 4B-C), and the correlation scatters plots indicate the correlation of the input variable and an immune cell/immunomodulator in a specific dataset (Figure 4D-E).

**Figure 4.**
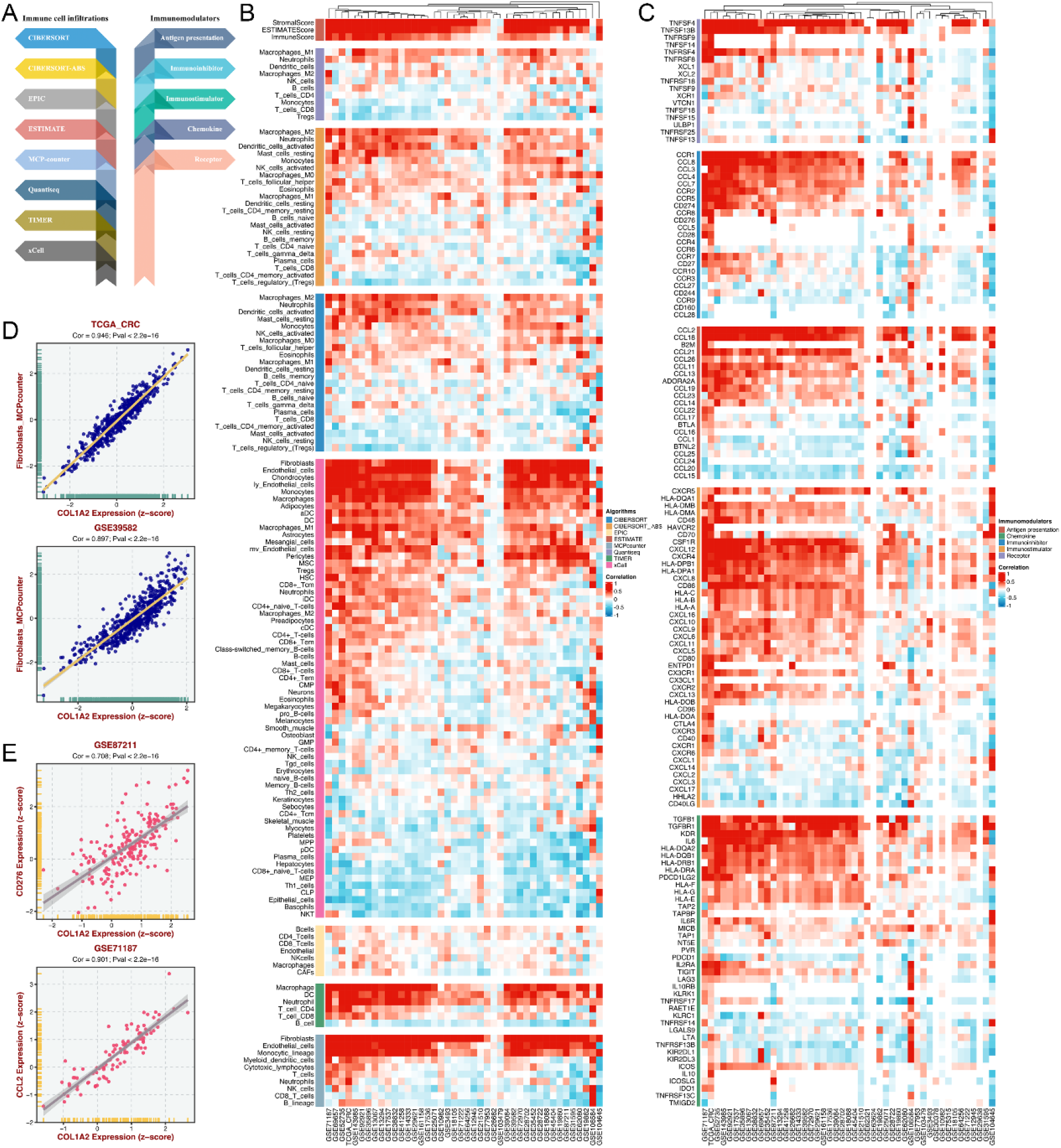
Modules for Cell infiltration and immunomodulator analysis. **A**. BEST offers eight immune infiltration assessment algorithms and five categories of immunomodulators. **B-C**. Heatmaps illustrate the correlations of the input variable with each immune cell (**B**) or immunomodulator (**C**) across all CRC datasets. **D-E**. Correlation scatters plots indicate the correlation of the input variable and an immune cell (**D**) or immunomodulator (**E**) in a specific dataset.

### Immunotherapy

To further investigate the clinical significance of the input variable in immunotherapies, we retrieved 19 immunotherapeutic cohorts with expression data and immunotherapy information (e.g., CAR-T, anti-PD-1, anti-CTLA4, etc.) (Figure 5A). Based on gene expression or gene set score in these datasets, users can conduct differential expression analysis (DEA) between response and non-response groups (Figure 5B), receiver operating characteristic (ROC) curve to evaluate the performance of the input variable in predicting the immunotherapeutic efficacy (Figure 5C), and survival analysis to assess the impact of the input variable on survival (OS and PFS) in immunotherapeutic cohorts that have undergone immunotherapies (Figure 5D).

**Figure 5.**
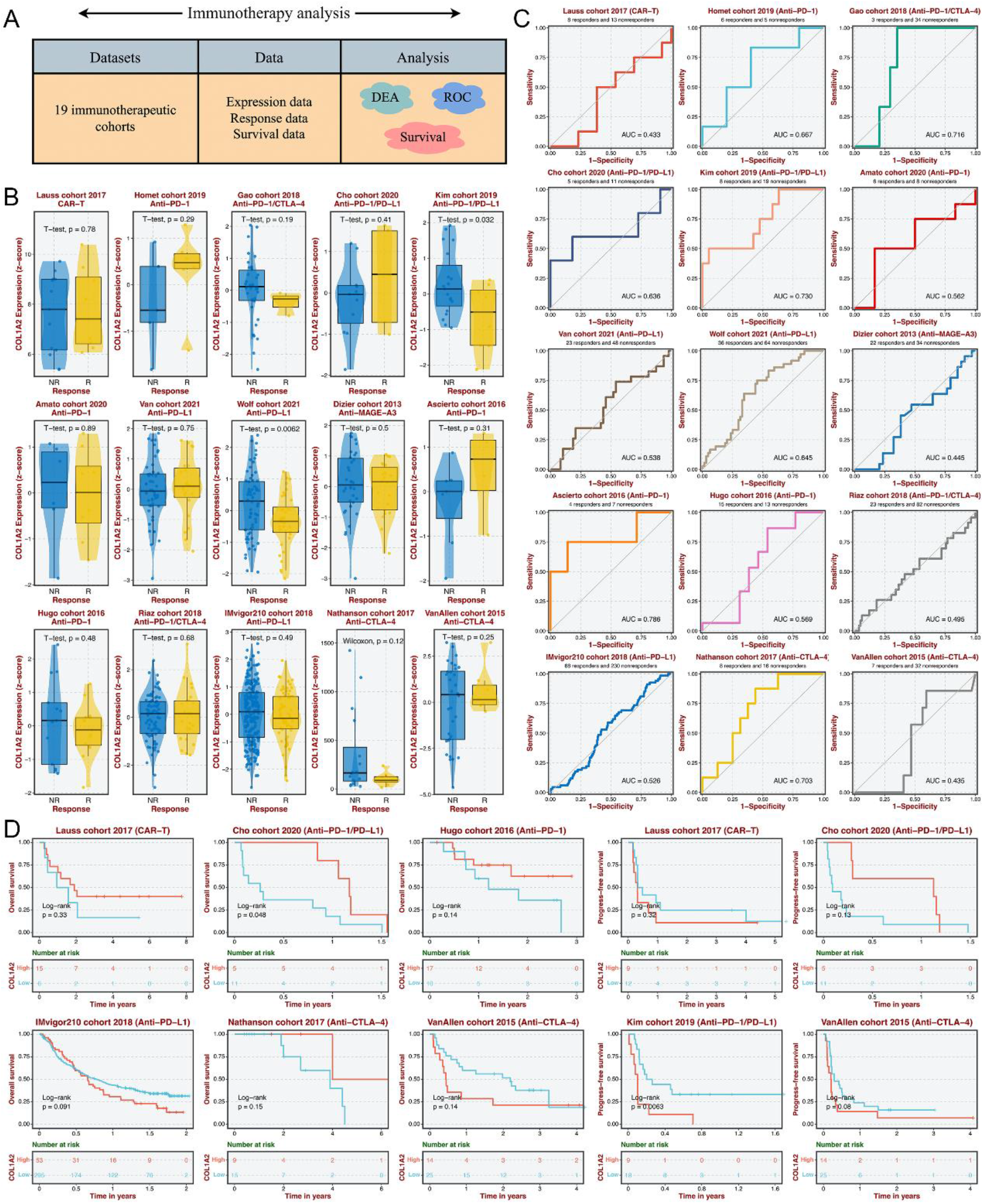
Immunotherapy analysis module. **A**. Schema describing data details and analysis for the immunotherapy module. **B**. Boxplots indicate the differential expression of *COL12A* between response and non-response groups. **C**. Receiver operating characteristic (ROC) curves evaluate the performance of the input variable in predicting the immunotherapeutic efficacy. **D**. Kaplan-Meier curves assess the impact of the input variable on survival (OS and PFS) in immunotherapeutic cohorts that have undergone immunotherapies.

### Candidate agents

In this analysis tab, BEST performs drug assessment in bulk samples based on drug responses and expression data of cancer cell lines from the GDSC_v1, GDSC_v2, CTRP, and PRISM databases (Figure 6A). Given the inherent differences between bulk samples and cell line cultures, we introduced a correlation of correlations framework^20^ to retain genes presenting analogical co-expression patterns in bulk samples and cell lines. As previously reported^21^, the model used for predicting drug response was the ridge regression algorithm implemented in the *oncoPredict* package^22^. This predictive model was trained on transcriptional expression profiles and drug response data of cancer cell lines with a satisfied predictive accuracy were evaluated by default 10-fold cross-validation, thus allowing the estimation of clinical drug response using only the expression data of bulk samples (Figure 6A). Modeling and prediction works have been completed, and drug assessments of all tumor samples based on four databases have been stored in the website background. BEST will calculate the correlations between all drugs and the input variable in all cohorts. According to the correlation rank of each drug across all datasets, we applied the robust rank aggregation (RRA)^23^ to determine drug importance related to the input variable (Figure 6B). Users can select the top drugs (self-defined number) to display the heatmap that illustrates the correlations of the input variable with each drug across all cohorts. Higher-ranked drugs indicate that high levels of the input variable predict drug resistance and vice versa. For example, high expression of *COL1A2* might suggest Afatinib resistance and Dasatinib sensitivity based on the GDSC_v2 database (Figure 6C). Also, users can select a drug database, a tumor dataset, and a specific drug to generate a correlation scatter plot (Figure 6D).

**Figure 6.**
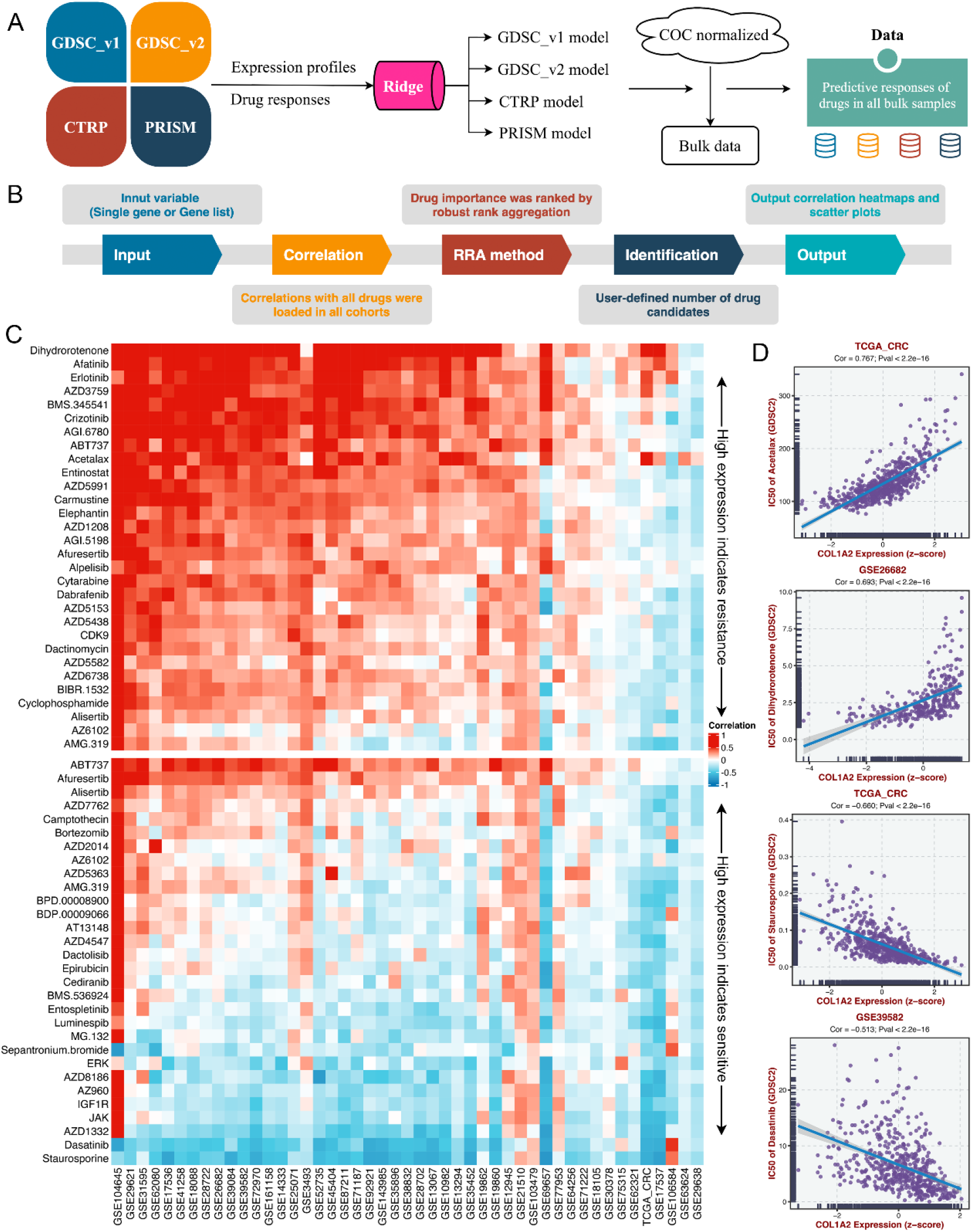
Candidate agents module. **A**. Overview of BEST performing drug assessment. **B**. Details of obtaining drugs that are significantly associated with the input variable. **C**. Heatmaps illustrate the correlations of the input variable with each drug across all cohorts. Higher-ranked drugs indicate that high levels of the input variable predict drug resistance and vice versa. **D**. Correlation scatters plots indicate the correlation of the input variable and a drug in a drug database and a tumor dataset.

### Genomic alteration

In this module, BEST has pre-processed mutation and copy number variation data from the TCGA database using *maftools*^24^ and *GISTIC2*.*0*^25^, respectively. Users can obtain a heatmap indicating genomic alterations as the input variable increase. The right panel of heatmap also displays the proportion of genomic alteration and statistical differences between the high and low groups. For example, with the rise in *COL1A2* expression, the genomics landscape of the TCGA-CRC dataset is illustrated in Figure 7. We found that the loss of chromosome segment 1p13.2 was more frequent in the high expression group.

**Figure 7.**
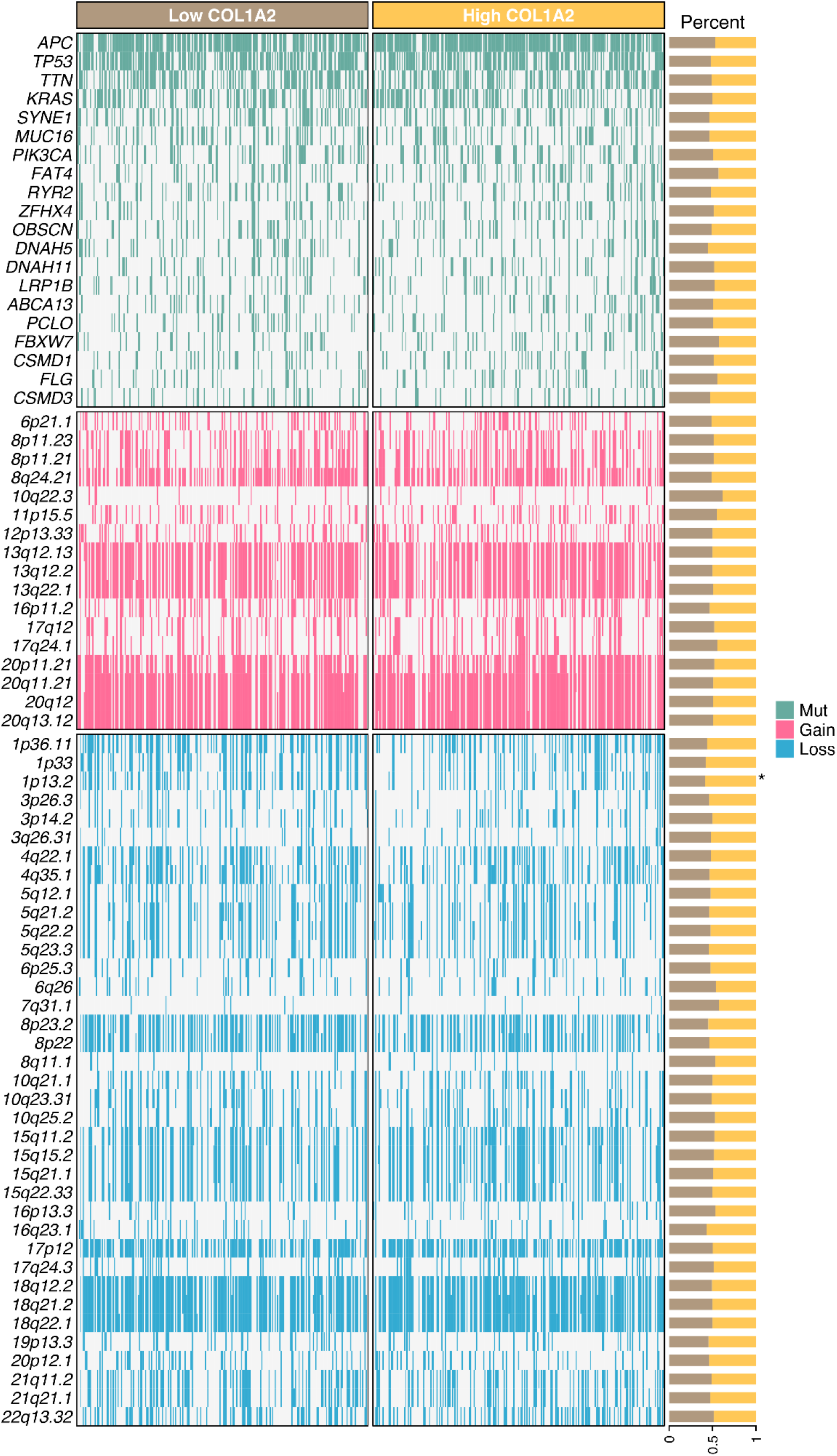
Genomic alteration analysis module.

## Discussion

As an interactive web tool, BEST aims to explore the clinical significance and biological functions of cancer biomarkers through large-scale data. Therefore, data richness is the foundation of BEST. From data collection, re-annotation, pre-processing, and pre-calculation to storage, we provide a tidy and uniform pan-cancer database, allowing users to call and interpret data quickly. BEST offers prevalent analysis modules to enable researchers without computational programming skills to conduct various bioinformatics analyses. Compared with other available tools^5–8, 26–28^, BEST has more datasets and more diverse analysis options, which complements well with them (Table 1).

**Table 1.**
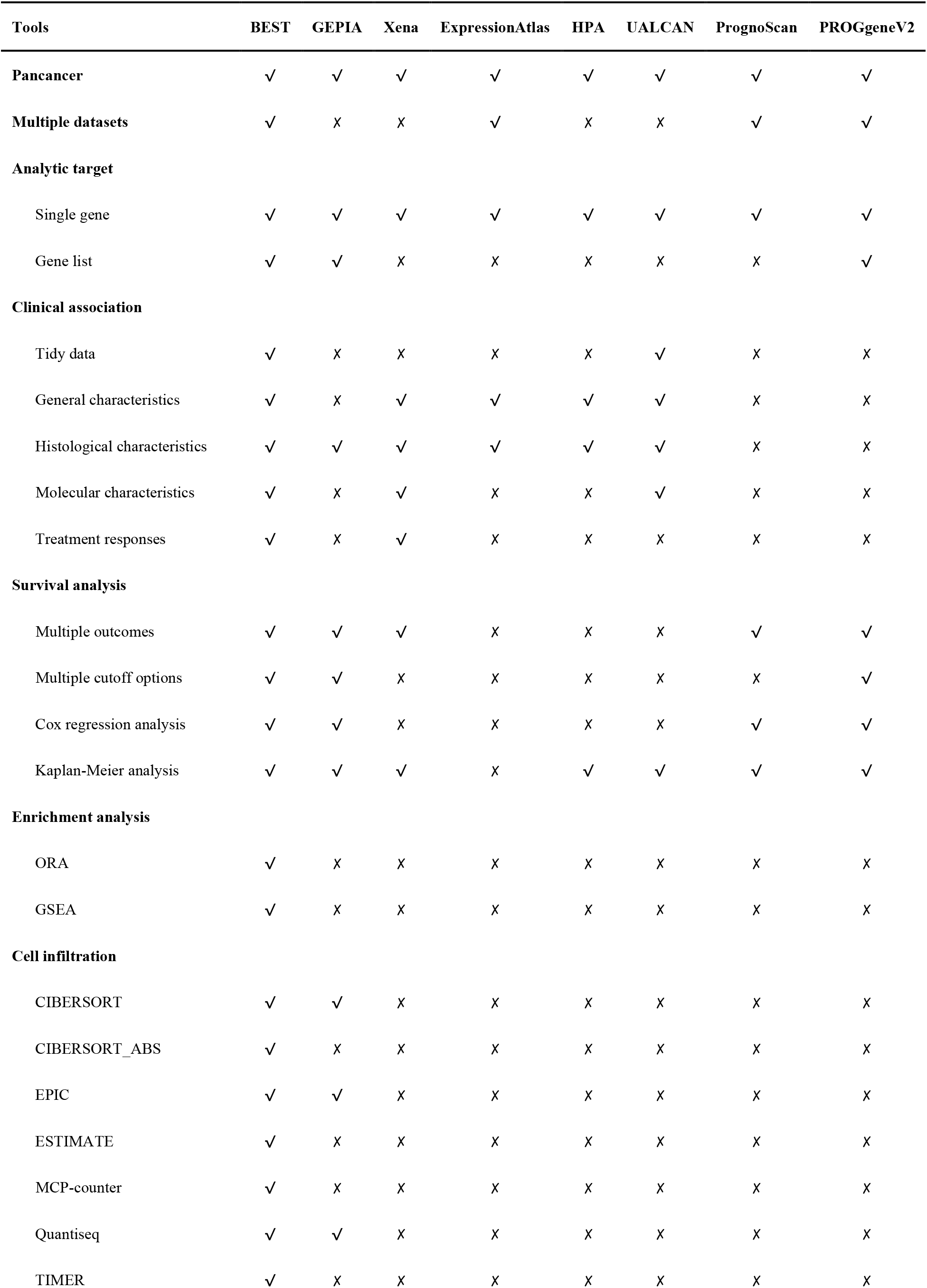
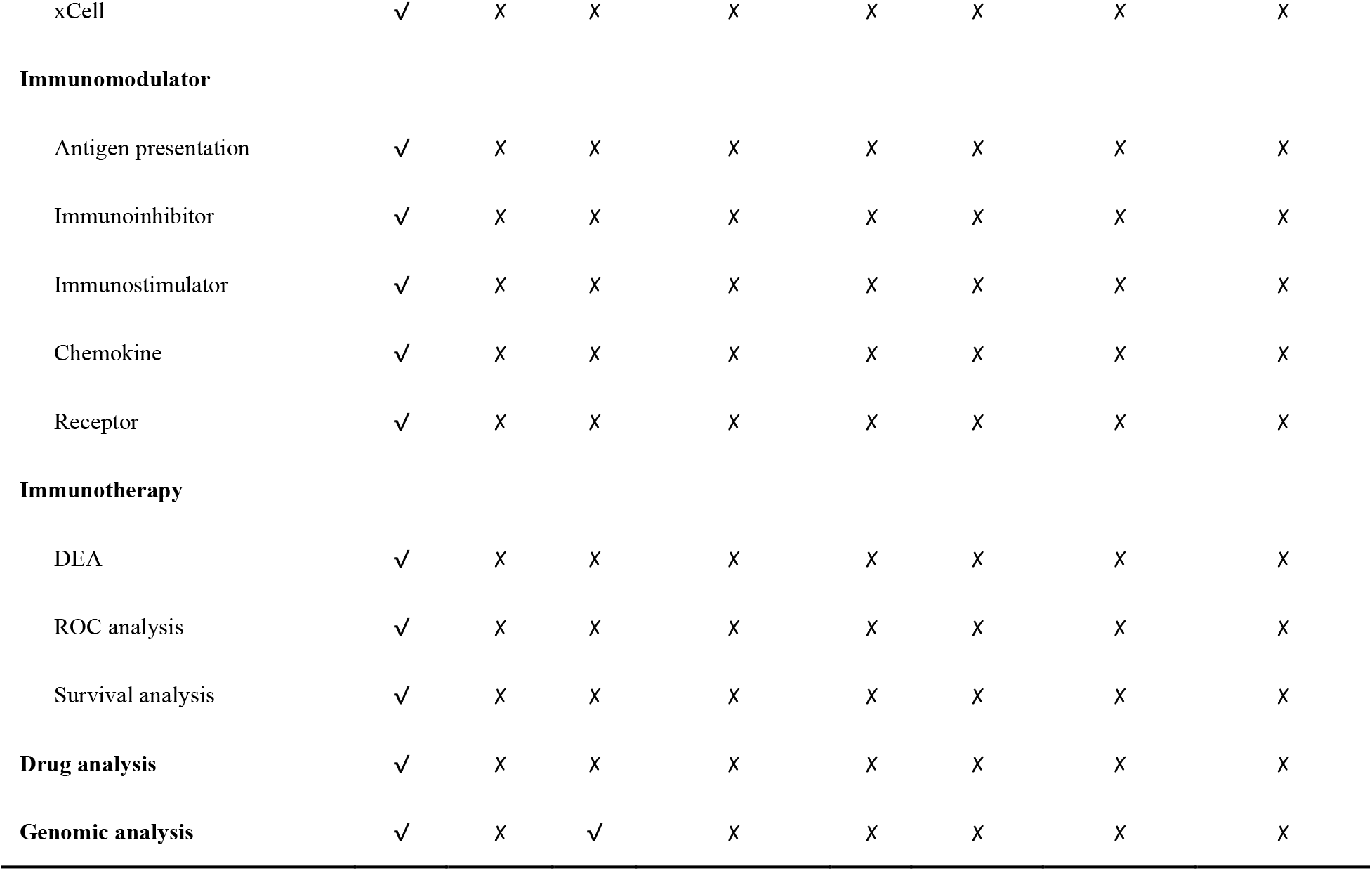
Comparison of BEST with other tools.

In BEST web application, users can identify cancer biomarkers associated with critical clinical traits (e.g., stage and grade), prognosis, and immunotherapy. Moreover, the underlying mechanisms of these biomarkers could be further explored using the enrichment, cell infiltration, and immunomodulator analysis modules. Users can also apply the candidate agent analysis tab to investigate high levels of cancer biomarkers that might indicate which drugs are resistant and which are sensitive to specific cancer.

Taken together, BEST provides a curated database and innovative analytical pipelines to explore cancer biomarkers at high resolution. It is an easy-to-use and time-saving web tool that allows users, especially clinicians and biologists without background knowledge of bioinformatics data mining, to comprehensively and systematically explore the clinical significance and biological function of cancer biomarkers. With constant user feedback and further improvement, BEST is promising to serve as an integral part of routine data analyses for researchers.

### Experimental procedures

### Data collection

BEST is committed to identifying robust tumor biomarkers through large-scale data. Hence, we retrieved cancer datasets with both expression data and important clinical information (e.g., survival, therapy, etc.) as much as possible. Eligible datasets were mainly enrolled from five databases, including The Cancer Genome Atlas Program (TCGA, https://portal.gdc.cancer.gov), Gene Expression Omnibus (GEO, https://www.ncbi.nlm.nih.gov/geo/), International Cancer Genome Consortium (ICGC, https://dcc.icgc.org), Chinese Glioma Genome Atlas (CGGA, http://www.cgga.org.cn/), and ArrayExpress (https://www.ebi.ac.uk/arrayexpress/). In total, we included more than 50,000 samples from 364 datasets for 27 cancer types.

### Data re-annotation and pre-processing

Raw expression data were extracted for subsequent processing (Figure 1). Data were re-annotated if the original probe sequences were available based on the GRCh38 patch 13 sequences reference from GENCODE (https://www.gencodegenes.org/). For RNA-seq data, raw count read was converted to transcripts per kilobase million (TPM) and further log-2 transformed. The raw microarray data from Affymetrix®, Illumina®, and Agilent® were processed using the *affy*^29^, *lumi*^30^, and *limma*^31^ packages, respectively. The normalized matrix files were directly downloaded for microarray data from other platforms. Gene expression was further transformed into z-score across patients in each dataset. To make it easier for users to interpret and present analysis results, we cleaned and unified the clinical traits. Take KRAS mutation as an example, GSE39084^32^ named it ‘kras.gene.mutation.status’, ‘mutation’ was labeled ‘M’ and ‘wild type’ was labeled ‘WT’; whereas GSE143985^33^ named it ‘kras_mutation’, ‘mutation’ was labeled ‘Y’ and ‘wild type’ was labeled ‘N’. We uniformly termed it ‘KRAS’, and ‘mutation’ was labeled ‘Mut’ and ‘wild type’ was labeled ‘WT’.

### Data calculation and storage

A tremendous amount of calculations are involved in BEST analysis, we thus have completed the time-consuming calculations in advance and used R.data for storage. Users can directly call these data, significantly reducing the user’s waiting time and background computing pressure. Take colorectal cancer (CRC) as an example, we collected a total of 47 datasets. Drug assessment is an analysis module of BEST, which requires fitting ridge regression models for individual drugs based on drug responses and expression data of cancer cell lines from the GDSC_v1, GDSC_v2, CTRP, and PRISM databases, and then predicting the sensitivity of each drug for CRC samples from all collected datasets. Apparently, if these results are not calculated in advance, users may have to wait more than three days. The pre-calculated content is displayed in Figure 1.

### Implementations

BEST is entirely free for users, built by the Shiny app and the HTML5, CSS, and JavaScript libraries for the client-side user interface. The Shiny app mainly executes data processing and analysis. The function of BEST is divided into eight tabs (Figure 1): Clinical association, Survival analysis, Enrichment analysis, Cell infiltration, Immunomodulator, Immunotherapy, Candidate agents, and Genomic alteration. Analysis results include images and tables, images can be downloaded in portable document format (PDF) and portable network graphics (PNG) format, and tables can be obtained in comma-separated value (CSV) format.

### Availability of data and materials

BEST is available at https://rookieutopia.com/.

## Funding or Acknowledgments

This study was supported by The Collaborative Innovation Major Project of Zhengzhou (Grant No. 20XTZX08017), The National Natural Science Foundation of China (Grant No. 82002433), and Science and Technology Project of Henan Provincial Department of Education (Grant No. 21A320036).

## Author contributions

ZQL, XWH, LL designed this work. ZQL, LL, SYW, and HX integrated and analyzed the data. ZQL, ZX, YQR, XYG, CGG, LBW, and LFL wrote this manuscript. ZQL, QC, PL, JZ, and XWH edited and revised the manuscript. All authors approved this manuscript.

## References

1. Hamilton, P. T., Anholt B. R., and Nelson B. H. (2022). Tumour immunotherapy: lessons from predator-prey theory. Nat Rev Immunol.https://doi.org/10.1038/s41577-022-00719-y.

2. Vesely, M. D., Zhang T., and Chen L. (2022). Resistance Mechanisms to Anti-PD Cancer Immunotherapy. Annu Rev Immunol. 40, 45–74.https://doi.org/10.1146/annurev-immunol-070621-030155.

3. Liu, Z., Liu L., Weng S., Guo C., Dang Q., Xu H., Wang L., Lu T., Zhang Y., Sun Z., et al. (2022). Machine learning-based integration develops an immune-derived lncRNA signature for improving outcomes in colorectal cancer. Nat Commun. 13, 816.https://doi.org/10.1038/s41467-022-28421-6.

4. Liu, Z., Guo C., Dang Q., Wang L., Liu L., Weng S., Xu H., Lu T., Sun Z., and Han X. (2022). Integrative analysis from multi-center studies identities a consensus machine learning-derived lncRNA signature for stage II/III colorectal cancer. EBioMedicine. 75, 103750.https://doi.org/10.1016/j.ebiom.2021.103750.

5. Tang, Z., Li C., Kang B., Gao G., Li C., and Zhang Z. (2017). GEPIA: a web server for cancer and normal gene expression profiling and interactive analyses. Nucleic Acids Res. 45, W98–W102.https://doi.org/10.1093/nar/gkx247.

6. Goldman, M. J., Craft B., Hastie M., Repecka K., McDade F., Kamath A., Banerjee A., Luo Y., Rogers D., Brooks A. N., et al. (2020). Visualizing and interpreting cancer genomics data via the Xena platform. Nat Biotechnol. 38, 675–678.https://doi.org/10.1038/s41587-020-0546-8.

7. Petryszak, R., Keays M., Tang Y. A., Fonseca N. A., Barrera E., Burdett T., Fullgrabe A., Fuentes A. M., Jupp S., Koskinen S., et al. (2016). Expression Atlas update--an integrated database of gene and protein expression in humans, animals and plants. Nucleic Acids Res. 44, D746–752.https://doi.org/10.1093/nar/gkv1045.

8. Uhlen, M., Fagerberg L., Hallstrom B. M., Lindskog C., Oksvold P., Mardinoglu A., Sivertsson A., Kampf C., Sjostedt E., Asplund A., et al. (2015). Proteomics. Tissue-based map of the human proteome. Science. 347, 1260419.https://doi.org/10.1126/science.1260419.

9. Hanzelmann, S., Castelo R., and Guinney J. (2013). GSVA: gene set variation analysis for microarray and RNA-seq data. BMC Bioinformatics. 14, 7.https://doi.org/10.1186/1471-2105-14-7.

10. Barbie, D. A., Tamayo P., Boehm J. S., Kim S. Y., Moody S. E., Dunn I. F., Schinzel C., Sandy P., Meylan E., Scholl C., et al. (2009). Systematic RNA interference reveals that oncogenic KRAS-driven cancers require TBK1. Nature. 462, 108–112.https://doi.org/10.1038/nature08460.

11. Lee, E., Chuang H. Y., Kim J. W., Ideker T., and Lee D. (2008). Inferring pathway activity toward precise disease classification. PLoS Comput Biol. 4, e1000217.https://doi.org/10.1371/journal.pcbi.1000217.

12. Tomfohr, J., Lu J., and Kepler T. B. (2005). Pathway level analysis of gene expression using singular value decomposition. BMC Bioinformatics. 6, 225.https://doi.org/10.1186/1471-2105-6-225.

13. Newman, A. M., Liu C. L., Green M. R., Gentles A. J., Feng W., Xu Y., Hoang C. D., Diehn M., and Alizadeh A. A. (2015). Robust enumeration of cell subsets from tissue expression profiles. Nat Methods. 12, 453–457.https://doi.org/10.1038/nmeth.3337.

14. Racle, J., de Jonge K., Baumgaertner P., Speiser D. E., and Gfeller D. (2017). Simultaneous enumeration of cancer and immune cell types from bulk tumor gene expression data. Elife. 6.https://doi.org/10.7554/eLife.26476.

15. Yoshihara, K., Shahmoradgoli M., Martinez E., Vegesna R., Kim H., Torres-Garcia W., Trevino V., Shen H., Laird P. W., Levine D. A., et al. (2013). Inferring tumour purity and stromal and immune cell admixture from expression data. Nat Commun. 4, 2612.https://doi.org/10.1038/ncomms3612.

16. Becht, E., Giraldo N. A., Lacroix L., Buttard B., Elarouci N., Petitprez F., Selves J., Laurent-Puig P., Sautes-Fridman C., Fridman W. H., et al. (2016). Estimating the population abundance of tissue-infiltrating immune and stromal cell populations using gene expression. Genome Biol. 17, 218.https://doi.org/10.1186/s13059-016-1070-5.

17. Finotello, F., Mayer C., Plattner C., Laschober G., Rieder D., Hackl H., Krogsdam, Loncova Z., Posch W., Wilflingseder D., et al. (2019). Molecular and pharmacological modulators of the tumor immune contexture revealed by deconvolution of RNA-seq data. Genome Med. 11, 34.https://doi.org/10.1186/s13073-019-0638-6.

18. Li, B., Severson E., Pignon J. C., Zhao H., Li T., Novak J., Jiang P., Shen H., Aster J. C., Rodig S., et al. (2016). Comprehensive analyses of tumor immunity: implications for cancer immunotherapy. Genome Biol. 17, 174.https://doi.org/10.1186/s13059-016-1028-7.

19. Aran, D., Hu Z., and Butte A. J. (2017). xCell: digitally portraying the tissue cellular heterogeneity landscape. Genome Biol. 18, 220.https://doi.org/10.1186/s13059-017-1349-1.

20. Guinney, J., Dienstmann R., Wang X., de Reynies A., Schlicker A., Soneson C., Marisa L., Roepman P., Nyamundanda G., Angelino P., et al. (2015). The consensus molecular subtypes of colorectal cancer. Nat Med. 21, 1350–1356.https://doi.org/10.1038/nm.3967.

21. Yang, C., Chen J., Li Y., Huang X., Liu Z., Wang J., Jiang H., Qin W., Lv Y., Wang H., et al. (2021). Exploring subclass-specific therapeutic agents for hepatocellular carcinoma by informatics-guided drug screen. Brief Bioinform. 22.https://doi.org/10.1093/bib/bbaa295.

22. Maeser, D., Gruener R. F., and Huang R. S. (2021). oncoPredict: an R package for predicting in vivo or cancer patient drug response and biomarkers from cell line screening data. Brief Bioinform. 22.https://doi.org/10.1093/bib/bbab260.

23. Kolde, R., Laur S., Adler P., and Vilo J. (2012). Robust rank aggregation for gene list integration and meta-analysis. Bioinformatics. 28, 573–580.https://doi.org/10.1093/bioinformatics/btr709.

24. Mayakonda, A., Lin D. C., Assenov Y., Plass C., and Koeffler H. P. (2018). Maftools: efficient and comprehensive analysis of somatic variants in cancer. Genome Res. 28, 1747–1756.https://doi.org/10.1101/gr.239244.118.

25. Mermel, C. H., Schumacher S. E., Hill B., Meyerson M. L., Beroukhim R., and Getz G. (2011). GISTIC2.0 facilitates sensitive and confident localization of the targets of focal somatic copy-number alteration in human cancers. Genome Biol. 12, R41.https://doi.org/10.1186/gb-2011-12-4-r41.

26. Chandrashekar, D. S., Bashel B., Balasubramanya S. A. H., Creighton C. J., Ponce-Rodriguez I., Chakravarthi B., and Varambally S. (2017). UALCAN: A Portal for Facilitating Tumor Subgroup Gene Expression and Survival Analyses. Neoplasia. 19, 649–658.https://doi.org/10.1016/j.neo.2017.05.002.

27. Mizuno, H., Kitada K., Nakai K., and Sarai A. (2009). PrognoScan: a new database for meta-analysis of the prognostic value of genes. BMC Med Genomics. 2, 18.https://doi.org/10.1186/1755-8794-2-18.

28. Goswami, C. P., and Nakshatri H. (2014). PROGgeneV2: enhancements on the existing database. BMC Cancer. 14, 970.https://doi.org/10.1186/1471-2407-14-970.

29. Gautier, L., Cope L., Bolstad B. M., and Irizarry R. A. (2004). affy--analysis of Affymetrix GeneChip data at the probe level. Bioinformatics. 20, 307–315.https://doi.org/10.1093/bioinformatics/btg405.

30. Du, P., Kibbe W. A., and Lin S. M. (2008). lumi: a pipeline for processing Illumina microarray. Bioinformatics. 24, 1547–1548.https://doi.org/10.1093/bioinformatics/btn224.

31. Ritchie, M. E., Phipson B., Wu D., Hu Y., Law C. W., Shi W., and Smyth G. K. (2015). limma powers differential expression analyses for RNA-sequencing and microarray studies. Nucleic Acids Res. 43, e47.https://doi.org/10.1093/nar/gkv007.

32. Kirzin, S., Marisa L., Guimbaud R., De Reynies A., Legrain M., Laurent-Puig P., Cordelier P., Pradere B., Bonnet D., Meggetto F., et al. (2014). Sporadic early-onset colorectal cancer is a specific sub-type of cancer: a morphological, molecular and genetics study. PLoS One. 9, e103159.https://doi.org/10.1371/journal.pone.0103159.

33. Shinto, E., Yoshida Y., Kajiwara Y., Okamoto K., Mochizuki S., Yamadera M., Shiraishi T., Nagata K., Tsuda H., Hase K., et al. (2020). Clinical Significance of a Gene Signature Generated from Tumor Budding Grade in Colon Cancer. Ann Surg Oncol. 27, 4044–4054.https://doi.org/10.1245/s10434-020-08498-3.

